# Divergent effects of cytomegalovirus and rheumatoid arthritis on senescent CD4^+^ T cells

**DOI:** 10.1101/2024.12.11.627966

**Authors:** Lea Williams, Ali O. Saber, Ruozhang Xu, Hannah Jung, Silina Awad, Anupama Shahane, Joshua F. Baker, Laura F. Su

## Abstract

CD4^+^ T cell senescence has been linked to repeated antigen stimulation. However, how different types of chronic antigenic exposures impact T cell differentiation and function remains incompletely understood. Using rheumatoid arthritis (RA) and cytomegalovirus (CMV) as models for persistent stimulation in autoimmunity and chronic viral infection, we performed high-dimensional mass cytometry analyses to examine their effects on CD4^+^ T cell differentiation. We found that CMV seropositive adults have a significant population of highly differentiated CD27^-^CD28^-^CD4^+^ T cells that exhibited common features of senescence, including CD57 and cytotoxic granule expression. In contrast, CD27^-^CD28^-^CD4^+^ T cells were rare in RA patients and Epstein-Barr virus (EBV) or Herpes simplex virus (HSV) seropositive individuals. The few that were present exhibited a predominantly non-cytotoxic profile, with higher expression of TCF1, CD127, and Ki67. Among CMV seropositive individuals, RA was associated with reduced degranulation of cytotoxic granules and lower cytokine production by senescent CD4^+^ T cells. We did not find an association with age, sex, clinical characteristics, or medication usage by univariate linear regression analyses. However, *in vitro* tofacitinib treatment reduced T cell functional activity, suggesting contributions from both the disease and RA treatment. These data uncovered distinct influences from CMV and RA, and their combined impact on senescent CD4^+^ T cell differentiation and function.

## Introduction

Chronic antigen stimulation drives terminal T cell differentiation. In patients with rheumatoid arthritis (RA), a highly differentiated CD4^+^ T cell population has been linked to increased disease activity, joint damage, and cardiovascular disease (1-5). These end-differentiated CD4^+^ T cells, exhibiting replicative senescence, frequently lacked costimulatory receptors and expressed CD57, a terminally sulfated carbohydrate epitope (6). They are highly pro-inflammatory, producing IFN-γ and TNF-α and expressing cytotoxic proteins, resembling cytotoxic CD8^+^ T cells (7).

Similar phenotypic subsets have been identified in other autoimmune diseases, infections, and cancer (8). For example, a cytotoxic CD27^-^CD28^-^CD4^+^ population emerges after primary Cytomegalovirus (CMV) infection and is expanded during viral latency (9, 10). An accumulation of replication-impaired CD57^+^CD4^+^ and CD8^+^ cells has also been observed in HIV infection (11, 12). In contrast to their presumed pathologic roles in autoimmunity, cytotoxic CD4^+^ T cells can be protective and have been shown in mice to provide direct defense against lethal influenza infection (13). In a human influenza challenge study, a higher baseline frequency of cytotoxic CD4^+^ T cells correlated with decreased viral shedding, lower symptom scores, and reduced disease duration after exposure (14). Recent studies on tumor-infiltrating T cells have also shown that cytotoxic CD4^+^ T cells play a critical role in tumor surveillance, with the ability to kill tumor cells in a class II MHC-dependent manner (15-18). In supercentenarians, these cells are significantly more abundant than in younger old adults and are thought to enhance anti-tumor and anti-viral immunity (19).

These studies highlight the various context-dependent roles of this unique CD4^+^ T cell subset. However, in the complex human environment, exposures seldom occur in isolation. How multiple co-existing stimuli converge to influence T cell baseline states remains unclear. Using high-dimensional mass cytometry, we analyzed CD4^+^ T cells from CMV seropositive and seronegative individuals, with and without RA, to investigate disease- and infection-related changes in T cell differentiation. Our data identified CMV as the primary driver of CD4^+^ T cell senescence. In the absence of CMV, RA patients and those with EBV or HSV infections had few CD27^-^ CD28^-^CD4^+^ T cells, which displayed minimal senescent features. However, RA modulated senescent CD4^+^ T cell function in CMV co-infected individuals, resulting in reduced degranulation and cytokine production. Univariate linear regression analyses did not identify a significant association between senescent cell function and clinical characteristics or medication usage, although drug-mediated effects likely contribute. *In vitro*, tofacitinib exposure reduced T cell responses to stimulation. Collectively, our findings underscore CMV’s critical influence on the composition of the human CD4^+^ T cell repertoire and reveal distinct effects of chronic viral infection and RA on senescent T cell differentiation and function.

## Results

### Senescent CD4^+^ T cells accumulate in CMV seropositive individuals

We performed mass cytometry with a 36-marker panel using PBMCs from 20 RA patients and 16 controls with similar age and sex, with or without a positive CMV IgG antibody test (Table 1). PBMCs were barcoded using Cell-ID, stained with metal-conjugated antibodies focusing on T cell differentiation, and acquired together on the mass cytometer. Data normalization across runs was performed using a bead-based standard (20). Live CD3^+^TCRαβ^+^CD4^+^ T cells were identified by manual gating and combined for analyses using the Spectre pipeline (Fig. S1A) (21). Non-linear dimensional reduction was performed using Uniform manifold approximation and projection (UMAP) (Fig. 1A). We observed two CD45RA^high^CD95^low^ naïve-like T cell clusters (clusters 5 and 7), a CXCR5^+^ T follicular helper cell-like cluster (cluster 8), and a CXCR3^+^ Th1-like cluster (cluster 4). Within memory cells, we observed a gradual loss of the co-stimulatory receptors, CD27 and CD28. Clusters 0 and 2 lacked CD27 but retained CD28, while cells negative for both CD27 and CD28 clustered in a distinct part of the UMAP (cluster 10). Cells expressing CD57 or CD45RA were present in this cluster, occupying both shared and separate areas (Fig. 1D). These data suggest that markers such as CD28-null, CD57, and CD45RA re-expression in the memory population identify overlapping but distinct populations of highly differentiated CD4^+^ T cells.

**Table 1:**
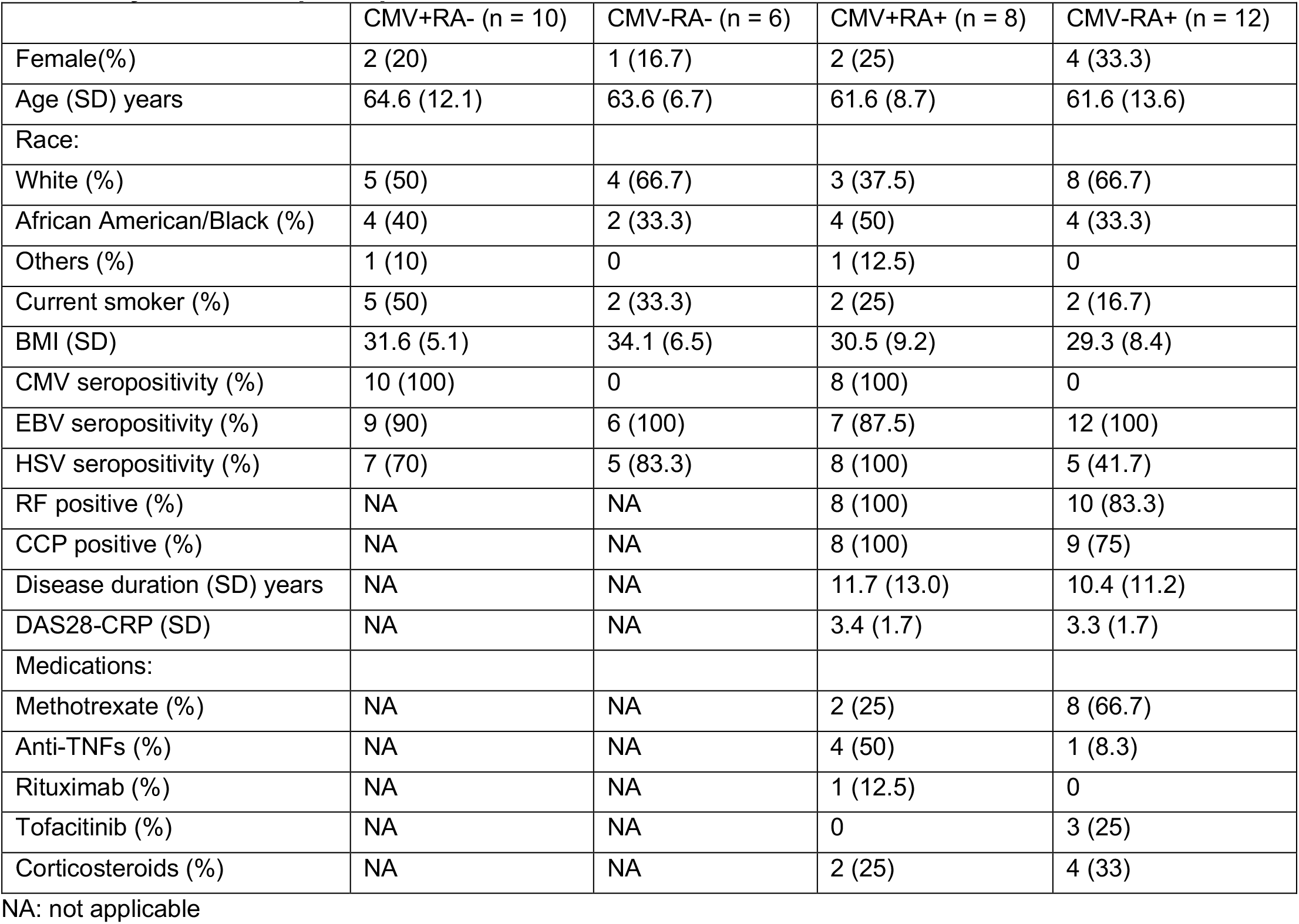
CyTOF cohort participant characteristics.

**Figure 1:**
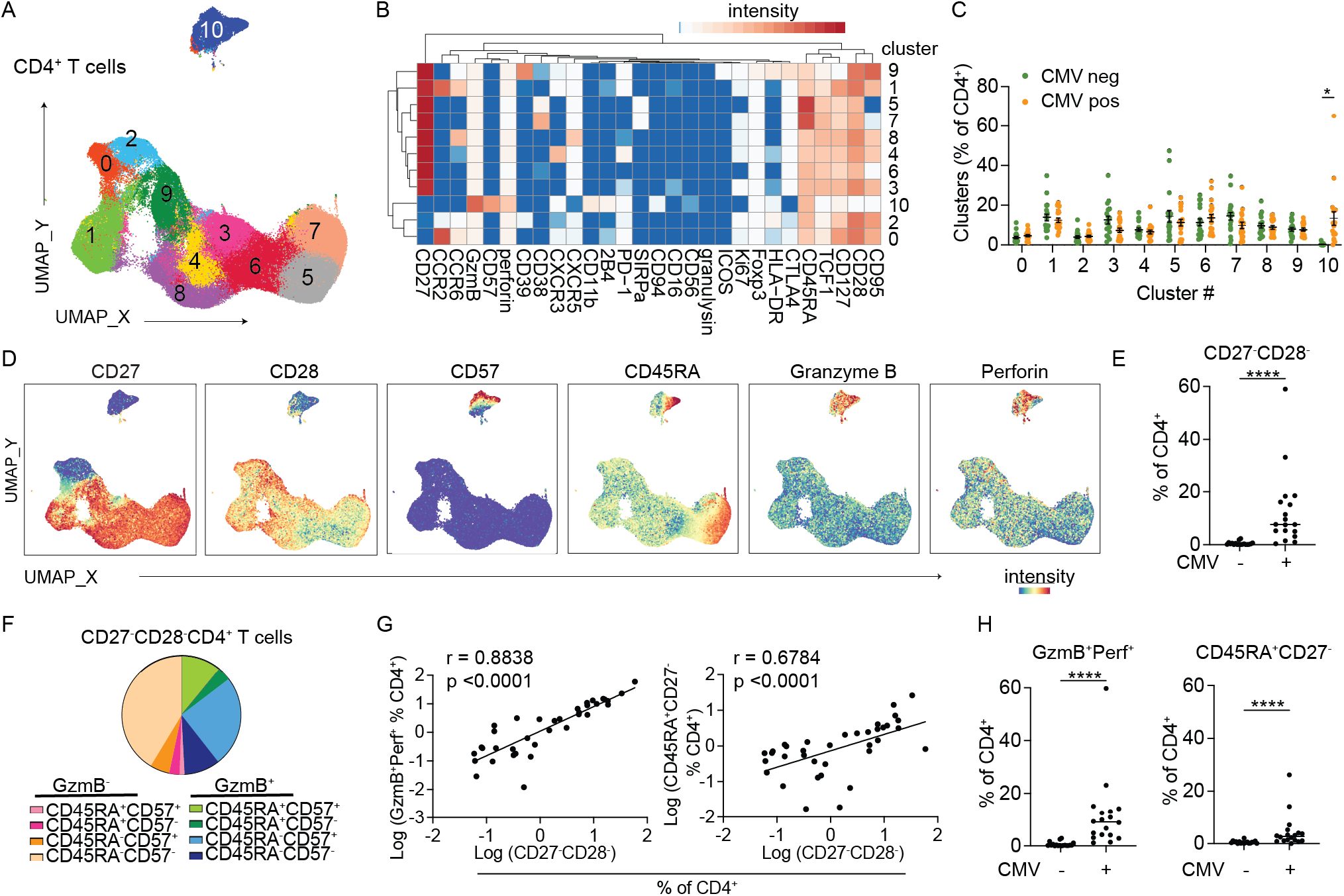
Cytotoxic end-differentiated CD4^+^ T cells are increased in CMV seropositive individuals. (A) UMAP displays Phenograph-defined clusters. Data combine 5,000 manually gated CD19^-^CD3^+^TCRab^+^CD4^+^ cells from each donor (n = 36 donors). (B) Heatmap shows the median staining signal of individual markers for clusters shown in A. Markers used to select input cells were excluded. (C) Plot summarizes the percentage of CD4^+^ T cells in each cluster, divided by CMV serostatus. (D) UMAPs display the staining intensity of the indicated markers on CD4^+^ T cells. (E) The frequency of manually gated CD27^-^CD28^-^ subset as a percentage of CD4^+^ T cells in CMV seronegative and seropositive donors. (F) Co-expression of GzmB, CD45RA, and CD57 in CD27^-^ CD28^-^ CD4^+^ T cells. Marker combinations were determined using Boolean operators on manually defined gates. (G)The relationship between CD27^-^CD28^-^ CD4^+^ T cell frequency and cytotoxic (left) or TEMRA (right) subsets. (H)The frequencies of manually defined GzmB^+^Perf^+^ (left) or CD45RA^+^CD27^-^ (right) cells as a percentage of CD4^+^ T cells in CMV seronegative and seropositive donors. Each filled circle represents cells from one individual. For (C) RM two-way ANOVA with Sidak’s multiple comparison test was performed. For (E) and (H), Mann-Whitney test was used. For (G), spearman correlation was performed.

We focused on cluster 10 because these cells were significantly increased in CMV seropositive individuals (Fig. 1C). Notably, the majority of cells in cluster 10 exhibited cytotoxic features by granzyme B (GzmB) and perforin expression (Fig. 1D). They also expressed higher levels of CD11b, 2B4, and CD94, which are more typically detected in NK cells and cytotoxic CD8^+^ T cells (Fig.1B). Similar to Phenograph defined cluster 10, the frequency of manually gated CD27^-^CD28^-^CD4^+^ T cells were increased in CMV seropositive individuals (Fig. 1E, S1A). On average, approximately half of the CD27^-^CD28^-^ subset expressed GzmB; of these, 73% expressed both GzmB and CD57, while 22% were positive for GzmB, CD57, and CD45RA. Within the GzmB negative subset, the majority were also negative for CD45RA and CD57 (Fig. 1F). Higher cytotoxic granule content and TEMRA differentiation positively correlated with CD27^-^CD28^-^CD4^+^ T cell expansion and were increased in CMV seropositive individuals (Fig1G-H, S1B-C).

To further evaluate the heterogeneity of CD27^-^CD28^-^CD4^+^ T cells, manually gated CD27^-^CD28^-^CD4^+^ T cells from both CMV seropositive and seronegative donors were exported and combined for spectral analyses. This showed a dominant CMV-associated difference in their differentiation states. While an average of 71% of CD27^-^ CD28^-^ subset from CMV seropositive individuals expressed GzmB and perforin, only 18% were GzmB^+^Perf^+^ in CD27^-^CD28^-^CD4^+^ T cells from people without CMV infection (Fig. 2A). Instead, CD27^-^CD28^-^CD4^+^ T cells from CMV seronegative individuals expressed markers of activation (HLA-DR), proliferation (Ki67), and self-renewal (TCF1, CD127) (Fig. 2B). On the UMAP, these cells clustered in a distinct region and were negative for GzmB and perforin expression (Fig. 2B-C). Manually defined Ki67^+^HLA-DR^+^ and TCF1^+^CD127^+^ subsets within CD27^-^ CD28^-^CD4^+^ T cells were correlated and more abundant in donors without CMV infection (Fig. 2D-F). Their frequencies showed a negative relationship with the size of the CD27^-^CD28^-^CD4^+^ T cell population, suggesting a loss of less-differentiated states during CD27^-^CD28^-^ T cell expansion (Fig. 2G). Thus, CD27^-^CD28^-^CD4^+^ T cells encompass multiple differentiation states, with features of senescence developing in a CMV-dependent manner.

**Figure 2:**
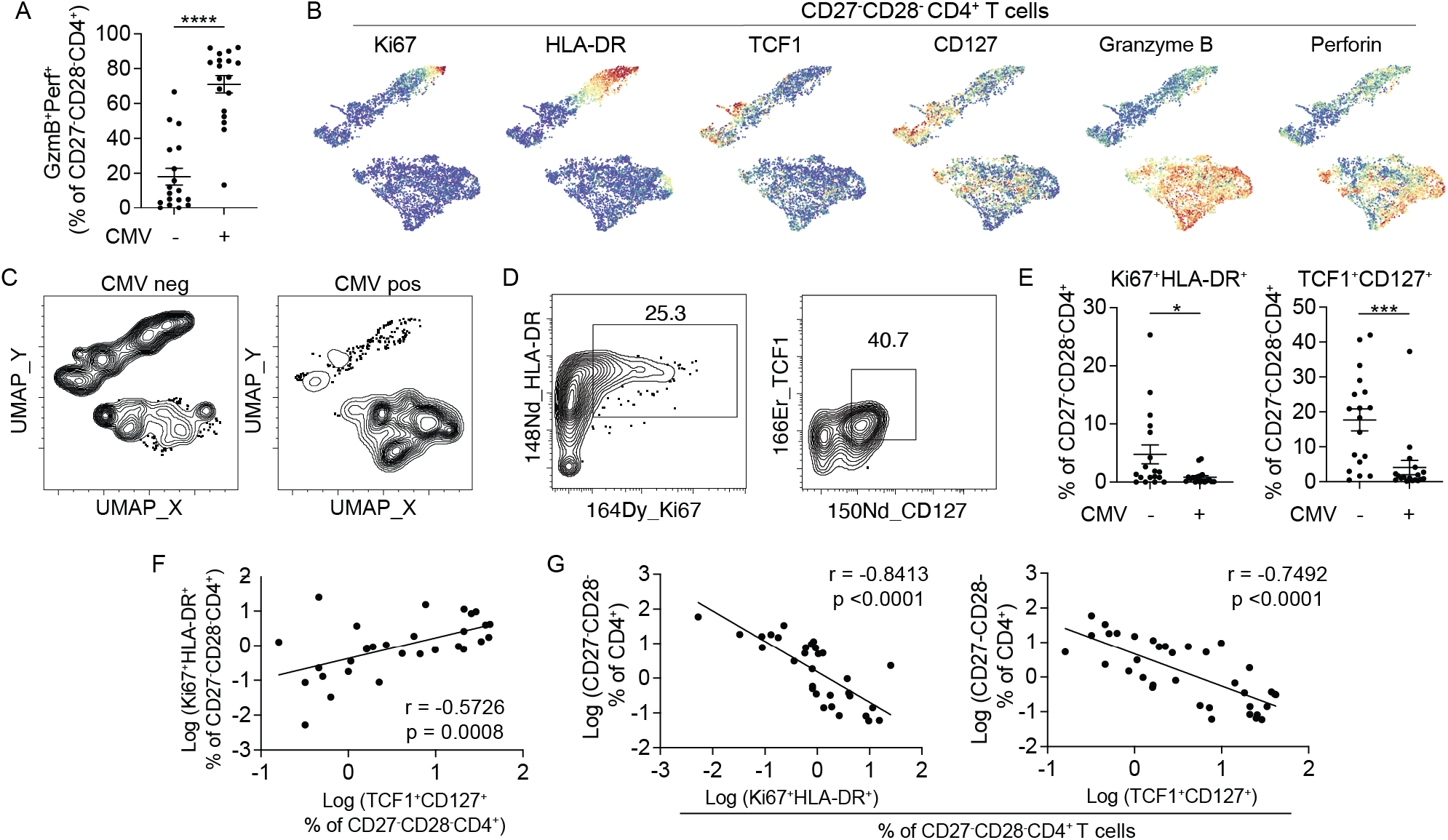
CD27^-^CD28^-^CD4^+^ T cells lack senescent features in CMV seronegative individuals. (A) The frequency of GzmB^+^Perf^+^ cells within the CD27^-^CD28^-^CD4^+^ subset in CMV seronegative and seropositive donors. (B) UMAPs display the staining intensity of the indicated markers. Data combine comparable numbers of cells from CMV seropositive and seronegative individuals for a total of 6016 manually gated CD27^-^CD28^-^CD4^+^ T cells. (C) UMAP distribution of CD27^-^CD28^-^CD4^+^ T cells, separated by CMV status. (D) Representative FACS plots show HLA-DR, Ki67 (left) and TCF1, CD27 (right) staining in CD27^-^CD28^-^CD4^+^ T cells from a CMV seronegative individual. (E) Plot summarizes the frequencies of Ki67^+^HLA-DR^+^ (left) and TCF1^+^CD127^+^ (right) T cells as a percentage of CD27-CD28-CD4^+^ T cells in CMV seronegative or seropositive donors. (F) The relationship between the frequencies of TCF1^+^CD127^+^ and Ki67^+^HLA-DR^+^ subsets as a percentage of CD27^-^ CD28^-^CD4^+^ T cells. (G)The correlation between Ki67^+^HLA-DR^+^ (left) or TCF1^+^CD127^+^ (right) subset and the total CD27^-^CD28^-^CD4^+^ T cell frequency. Each filled circle represents data from one individual. For (A) and (E), Mann-Whitney test was used. Spearman correlation was performed for (F) and (G).

### CMV exerts a dominant effect on CD4^+^ T cell senescence over RA, EBV, and HSV

Next, we asked if RA independently contributes to the accumulation of senescent CD4^+^ T cells. To examine this, we divided CMV seropositive and negative individuals according to their RA status and generated UMAPs on CD27^-^CD28^-^CD4^+^ T cells from each group. This showed that cluster distribution was primarily influenced by CMV status, with similar patterns observed between individuals with or without RA (Fig. 3A). The frequency of cluster 10 from Spectre analyses, as well as CD27^-^CD28^-^ and GzmB^+^Perf^+^ CD4^+^ T cells identified by manual gating, also differed based on CMV but not RA status (Fig. 3B-D).

**Figure 3:**
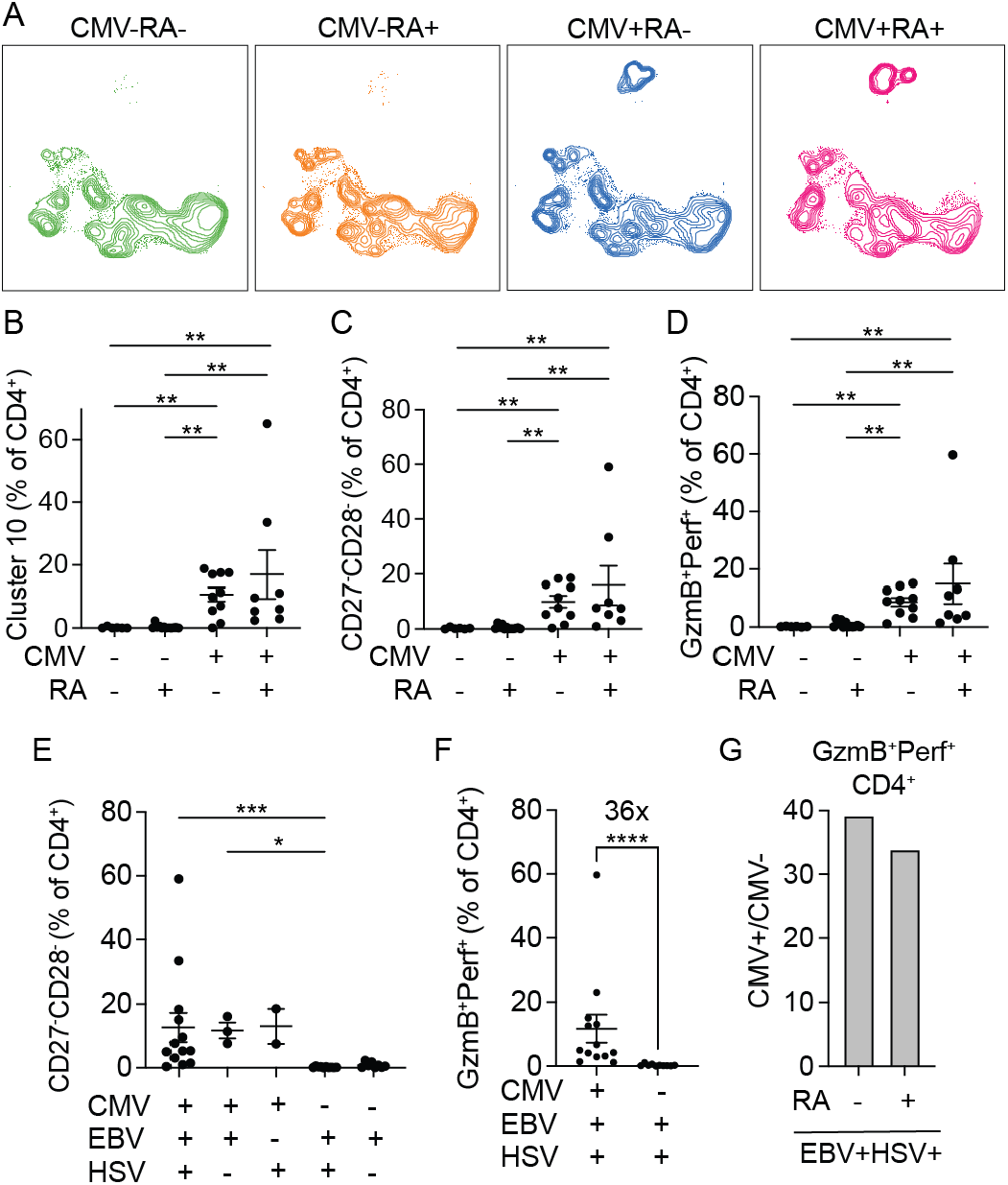
Senescent CD4^+^ T cell frequency depends on CMV and not RA, EBV, or HSV. (A) UMAPs of CD27^-^CD28^-^CD4^+^ T cells, separated by CMV serostatus in donors with or without RA. CMV-RA- (CMV seronegative controls), CMV-RA+ (CMV seronegative RA patients), CMV+RA- (CMV seropositive controls), CMV+RA+ (CMV seropositive RA patients). (B-D) Plots summarize the percentage of CD4^+^ T cells in cluster 10 (B), negative for CD27 and CD28 (C), or expressing GzmB and perforin (D) by CMV and RA status. (E) The frequencies of CD27^-^CD28^-^ subset as a percentage of CD4^+^ T cells, grouped by CMV, EBV, and HSV seropositivity. (F) The frequencies and the fold-difference of CD4^+^ T cells expressing GzmB and perforin in EBV and HSV positive donors, with or without CMV co-infection. (G) The ratio of averaged GzmB^+^Perf^+^ expression in CMV seropositive and seronegative individuals with EBV and HSV co-infection, grouped by RA status. Each filled circle represents data from one individual. (B-E) Kruskal-Wallis with Dunn’s multiple comparison tests were used. For (F), Mann-Whitney test was performed.

These data point to a dominant effect from CMV on CD4^+^ T cell terminal differentiation. Besides CMV, other latent viruses, including EBV and HSV, are also highly prevalent and infect the majority of the adult human population (22, 23). To compare the effects of different latent viral infections, serologic typing for EBV and HSV was performed and the data were grouped based on the serostatus for each virus. This showed that CD27^-^CD28^-^ CD4^+^ T cell frequency remains low despite seropositivity for EBV and/or HSV in the absence of CMV co-infection (Fig. 3E). Similarly, cytotoxic CD4^+^ T cells were infrequent in EBV and HSV seropositive individuals without CMV. In individuals positive for both EBV and HSV, CMV co-infection was associated with a 36-fold increase in GzmB^+^Perf^+^ CD4^+^ T cells, with similar levels observed in both controls and RA patients (Fig. 3F-G). Taken together, these data suggest that latent CMV infection disproportionally enhances senescent T cell differentiation, surpassing the influences of EBV, HSV, and RA.

### Senescent CD4^+^ T cells show impaired function in RA patients

Although we did not find a higher frequency of senescent CD27^-^CD28^-^CD4^+^ T cells in RA patients, RA and other disease-associated factors may influence their function. To test this, we recruited additional donors for a total of 24 controls and 24 RA patients, this time including only CMV seropositive individuals to focus on senescent CD4^+^ T cells (Table 2). PBMCs were stimulated with Staphylococcal Enterotoxin B (SEB) for 6 hours and analyzed for IFN-γ, TNF-α, and the degranulation marker, CD107a. Compared to controls, CD27^-^CD28^-^CD4^+^ T cells from RA patients showed diminished function, producing less pro-inflammatory cytokines TNF-α and IFN-γ (Fig. 4A-B, S2A). Cells from RA patients also showed reduced CD107a expression with unchanged granzyme B level, suggesting decreased degranulation capacity in cytotoxic cells (Fig. 4C, S2C-D). Overall, polyfunctional cells co-expressing TNF-α, IFN-γ, and CD107a were reduced in RA patients (Fig. 4D). In contrast, CD8^+^ T cell function was unaffected. CD27^-^CD28^-^CD8^+^ T cells from RA patients and controls produced similar levels of TNF-α and INF-γ after SEB stimulation. The proportion of CD107a^+^ and polyfunctional TNF-α^+^IFN-γ ^+^CD107^+^ cells within the CD27^-^CD28^-^CD8^+^ subset also did not differ significantly between individuals with or without RA (Fig. 4E-H).

**Table 2:** FACS cohort participant characteristics.

|  | Control (n = 24) | RA (n = 24) |
| --- | --- | --- |
| Female (%) | 12 (50.0) | 19 (79.2) |
| Age (SD) years | 43.5 (17.4) | 50.9 (16.8) |
| Race: |  |  |
| White (%) | 10 (41.7) | 8 (33.3) |
| African American/Black (%) | 5 (20.8) | 12 (50.0) |
| Asian (%) | 3 (12.5) | 0 (0) |
| Others (%) | 6 (25.0) | 4 (16.7) |
| Current smoker (%) | 5 (20.8) | 5 (20.8) |
| BMI (SD) | 33.6 (11.3) | 28.8 (6.2) |
| CMV seropositivity (%) | 24 (100) | 24 (100) |
| RF positive (%) | NA | 19 (79.2) |
| CCP positive (%) | NA | 16 (66.7) |
| Disease duration (SD) years | NA | 12.7 (23.5) |
| DAS28-CRP (SD) | NA | 2.25 (1.0) |
| Medications: |  |  |
| Methotrexate (%) | NA | 11 (45.8) |
| Anti-TNFs (%) | NA | 2 (8.4) |
| Tocilizumab (%) | NA | 1 (4.2) |
| Rituximab (%) | NA | 1 (4.2) |
| Tofacitinib (%) | NA | 1 (4.2) |
| Corticosteroids (%) | NA | 6 (25) |

NA: not applicable

**Figure 4:**
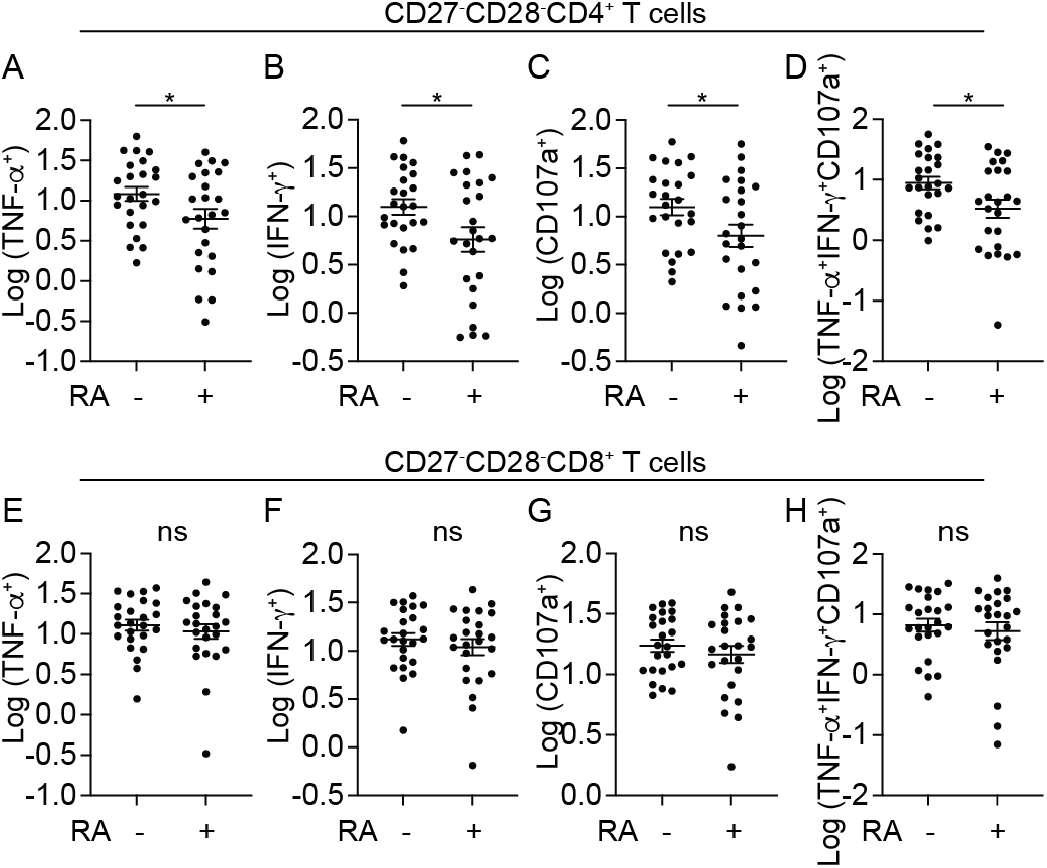
Senescent CD4^+^ T cells display decreased functional potential in RA patients. PBMCs from 24 controls and 24 RA patients were stimulated with SEB for 6 hours and analyzed for cytokine production and degranulation. Plot summarizes the frequencies of CD27^-^CD28^-^ T cells stained for TNF-α (A, E), IFN-γ (B, F), CD107 (C, G) or triple positive for TNF-α, IFN-γ, and CD107 (D, H) as a percentage of CD4^+^ or CD8^+^ T cells in controls and RA patients. Each filled circle represents data from one individual. Welch’s t-test was used.

Univariate linear regression analyses were performed to evaluate the relationship between cellular response and participant characteristics. We found no significant association between TNF-α^+^, IFN-γ ^+^, CD107a^+^, and TNF-α^+^IFN-γ ^+^CD107^+^ frequencies with age, sex, body mass index (BMI), smoking status, disease duration, c-reactive protein (CRP) levels, or disease activity scores as determined by DAS28-CRP. There was also no significant association with the use of methotrexate, prednisone, or either TNF inhibitor (TNFi) or non-TNFi biologic therapies after adjusting for age and sex in RA patients (Table S1). The lack of medication effect suggests that RA itself may directly contribute to the reduction in CD27^-^CD28^-^CD4^+^ T cell response. Consistent with this, two newly diagnosed RA patients who had not yet started treatment exhibited some of the lowest TNF-α^+^IFN-γ ^+^CD107^+^ T cell levels (Fig. S3). Alternatively, the lack of a significant medication association may reflect heterogeneous drug effects in a limited sample size. Among RA patients with a nearly undetectable TNF-α^+^IFN-γ ^+^CD107^+^ level in CD27^-^CD28^-^CD4^+^ T cells (<1%), one was taking a JAK inhibitor, tofacitinib, which is known to suppress T cell function (Fig. S3) (24). We further explored the potential effect of tofacitinib on CD27^-^CD28^-^CD4^+^ T cells by treating PBMCs *in vitro* with tofacitinib from 10 RA patients and 10 controls, selected based on sample availability. To examine tofacitinib’s effect on T cell basal state without directly impacting T cell activation (24, 25), we pretreated PBMCs with tofacitinib for 40 hours, removed it from the culture, followed by SEB stimulation and antibody staining. This showed reduced IFN-γ, TNF-α, CD107a expression, along with a decrease in polyfunctional TNF-α^+^IFN-γ ^+^CD107^+^ population in tofacitinib pre-treated cells (Fig. 5 A-D). Both CD4^+^ and CD8^+^ subsets were impacted, with inhibition being more pronounced for CD4^+^ T cells (Fig. 5E-I). T cells from RA patients and controls were similarly susceptible to tofacitinib treatment (Fig. 5J). Thus, tofacitinib exposure modifies the T cell baseline, impacting their subsequent activity *in vitro*. Collectively, our findings demonstrate a reduced functional capacity of senescent CD4^+^ T cells in RA patients, likely influenced by both the disease and RA treatment.

**Figure 5:**
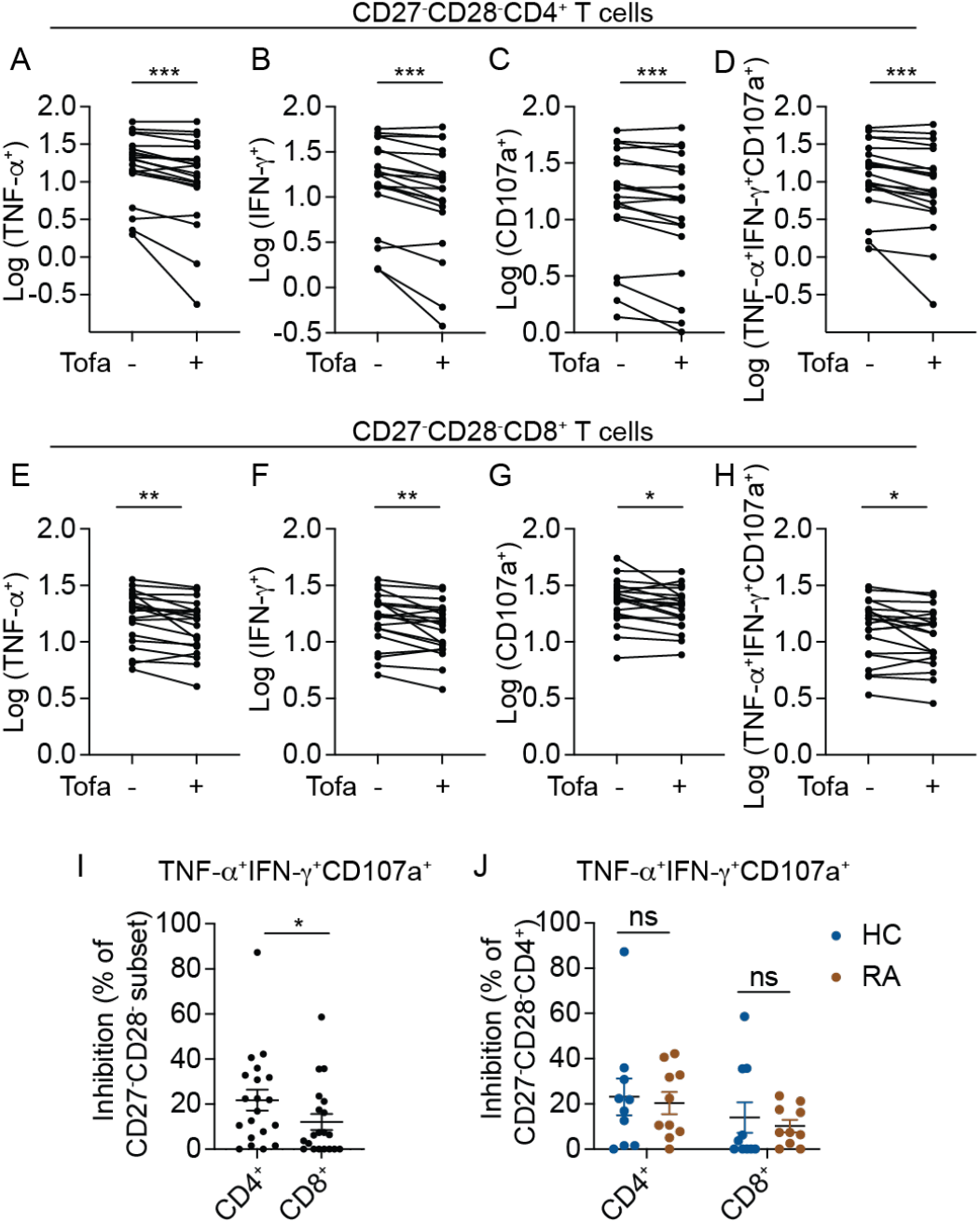
Tofacitinib inhibits the activities of CD27^-^CD28^-^ T cells *in vitro*. (A-H) PBMCs were incubated with or without 10uM tofacitinib for 40 hours, then replenished with fresh media and stimulated with SEB for 6 hours. Plots summarize the frequencies of the indicated subsets as a percentage of CD27^-^CD28^-^CD4^+^ T cells (A-D) or CD27^-^CD28^-^CD8^+^ T cells (E-H). (I-J) Tofacitinib inhibition of TNF-α^+^IFN-γ ^+^CD107^+^ response in CD27^-^ CD28^-^ CD4^+^ and CD27^-^CD28^-^ CD8^+^ T cells (I), separated by disease status (J). Inhibition was calculated as 1- (treated/untreated), with 0 as the lower bound. For (A), (D), (I), Wilcoxon matched-pairs signed rank test was performed. For (B-C and E-H), paired t-test was used. (J) uses RM two-way ANOVA.

## Discussion

Many RA patients are co-infected with CMV, a virus that infects approximately 50% of adults in the United States (26-28). Here, we performed high-dimensional phenotypic analyses on a cohort of RA patients and controls with known CMV serostatus to determine the individual contributions of CMV and RA to CD4^+^ T cell senescence. We began by analyzing the relationships between markers commonly used to identify late-differentiated CD4^+^ T cells. This showed that CD57 labeled a narrower subset of CD28-null cells, partially overlapping with TEMRA cells that re-expressed CD45RA. Cytotoxic features within the CD27^-^CD28^-^ subset were primarily observed in donors exposed to CMV. In the absence of latent CMV infection, CD27^-^CD28^-^ CD4^+^ T cells were minimally expanded. They also retained a less differentiated state, as suggested by TCF1, CD127, and Ki67 expression. RA and other latent viral infections we examined, including EBV and HSV, were insufficient to independently drive a significant expansion of senescent CD4^+^ T cells without CMV co-infection. This result differs from previous studies that reported an accumulation of senescent CD4^+^ T cells in RA patients (1-5). Selecting different marker combinations to define the senescent population did not change our findings. We speculate that the difference may be due to unaccounted effects from CMV. Consistent with this, previous studies have shown that CD28-null CD4^+^ T cells are significantly more abundant in CMV+ than in CMV-RA patients, although a direct comparison with non-RA controls was not available (29, 30). In addition, other differences in the selection of study participants may contribute. As most prior studies were conducted before or during the early years of biologic treatment, the impact of RA-mediated inflammation could have evolved as disease control improves over time.

While senescent CD4^+^ T cells were not more abundant in RA patients after accounting for CMV status, RA was associated with modified functional responses following T cell stimulation. Senescent CD4^+^ T cells in CMV seropositive RA patients secreted less IFN-γ and TNF-α and had reduced cytotoxic degranulation after stimulation compared to those from CMV seropositive individuals without RA. RA itself likely contributes to these findings. A highly impaired functional response could already be observed in two not yet treated RA patients in our cohort. While we did not identify a statistically significant association with medication usage, *in vitro* drug treatment with tofacitinib suggests at least the theoretical possibility that T cell inhibition from medications could reduce senescent T cell responses. Cytotoxic degranulation and cytokine production were more inhibited in senescent CD4^+^ T cells than CD27^-^CD28^-^CD8^+^ T cells in RA patients and following tofacitinib treatment in culture, suggesting a unique vulnerability in the CD4^+^ subset. Our data is limited to associations and does not assess the functional impact of these changes. However, given the growing recognition of cytotoxic CD4^+^ T cells in tumor surveillance and control, we speculate that inhibiting senescent CD4^+^ T cells, the main cytotoxic CD4^+^ population, may contribute to the increased cancer risk associated with tofacitinib in RA patients (31, 32). The impaired ability to eliminate virus-infected cells may also lead to increased infection and varicella-zoster virus reactivation in patients on a JAK inhibitor (33-35). Future studies will be needed to clarify the relative effects of RA and medications on these highly differentiated CD4^+^ T cells, their mechanisms of action, and functional implications.

In conclusion, CMV has a dominant effect on the accumulation of senescent CD4^+^ T cells. RA further modifies these cells and is associated with an impaired functional phenotype with reduced cytotoxic degranulation and cytokine production. Our study highlights the need to consider CMV and the broader history of exposures in future research on RA and other autoimmune diseases.

## Acknowledgments

We thank the study subjects for their participation.

## Author contributions

Conceptualization, L.F.S.; Experimentation, L.W., A.O.S. L.F.S; High-dimensional phenotypic analyses, R.X, A.O.S.; Study recruitment, H.J., S.A., A.S.; Modeling and statistical support, J.F.B.; Supervision, L.F.S.; Manuscript preparation, L.W., A.O.S., J.F.B., L.F.S.

## Methods

### Sex as a biological variable

Both male and female participants were included in this study. Sex was analyzed as a biological variable in the multiparameter linear regression analyses.

### Human samples

Blood samples were collected from healthy volunteers and rheumatology clinic patients at the University of Pennsylvania and Corporal Michael J Crescenz VA Medical Center from 2015 to 2023. All samples were de-identified and obtained with IRB regulatory approval from the University of Pennsylvania or Corporal Michael J Crescenz VA Medical Center, depending on the recruitment site. For sample processing, vacutainer tubes were spun down for plasma collection. PBMCs were isolated by density gradient centrifugation (Ficoll-Paque, GE Healthcare) and cryopreserved in fetal bovine serum (FBS) with 10% DMSO. Subject characteristics are shown in Tables 1 and 2.

### CMV, HSV, and EBV serologic testing

CMV, HSV, and EBV ELISA on plasma samples were performed and interpreted according to manufacturer instructions. Manufacturer-defined positivity index was used. Samples with ambiguous results were retyped and excluded if the result remained equivocal after 3 repeats. CMV typing was performed using Cytomegalovirus IgG ELISA Kit (KA1452, Abnova). HSV typing was performed using Simplex Virus I IgG ELISA Kit (KA0229, Abnova). EBV typing was performed using Epstein Barr Virus EBNA-1 IgG ELISA Kit (KA1448, Abnova).

### CyTOF staining and analyses

#### CyTOF staining

Cryopreserved PBMCs were thawed, washed, and incubated anti-CD127, anti-CCR2, anti-CD27, anti-CXCR3, anti-CCR6, and anti-TCRαβ for 30 minutes at room temperature before cisplatin staining, fixation, and cell ID barcoding according manufacturer protocol (StandardBioTools). Metal barcoded cells were pooled into a single tube and stained with the remainder of surface antibodies for 30 minutes at room temperature. For intracellular staining, cells were permeabilized and fixed using Foxp3 staining buffer set (eBioscience) and incubated with the intracellular antibody cocktail for 1 hour at room temperature (Table S2). Metal conjugation of CyTOF antibodies was performed according to the manufacturer protocol using the X8 Maxpar kit (StandardBioTools). After staining, cells were washed three times then resuspended in 2% paraformaldehyde (Electron Microscopy Sciences) with 125nM iridium intercalator (StandardBioTools) for an overnight incubation at 4°C. The next day, cells were washed three times, including a final wash in distilled water, and resuspended in water containing normalization beads before acquisition on Helios. Bead standards were used to normalize CyTOF runs with the Matlab-based Nolan lab normalizer (20).

#### Data analyses

Doublets and beads were excluded from Iridium^+^Cisplatin^-^ cells. From each sample, equal numbers of manually gated CD19^-^CD3^+^TCRαβ^+^CD4^+^ cells were downsampled, and exported using FlowJo (BD Biosciences). For CD4^+^ T cell analyses, a total of 180,000 cells were read into R by flowCore and combined into one single dataset for subsequent data processing and high-dimensional analyses using the Spectre package in R (36). Staining intensities were converted using Arcsinh transformation with a cofactor of 5. Clustering was performed using Phenograph with nearest neighbors set to 1100 (k = 1100) (37). Unbiased uniform manifold approximation and projection (UMAP) was used for dimensional reduction and visualization (38). Color scale was modified to use the same color for 0 and values under 0 after Arcsinh transformation. The heatmap was generated using the “gplot” package in R and showed the raw staining intensity of each marker after arcsinh transformation. Non-T cell markers and those used to select input cells were excluded. Heatmap dendrograms were clustered by Euclidean distance. CD27^-^CD28^-^CD4^+^ T cell analyses was performed as above using 6016 manually gated CD27^-^CD28^-^ population with a cofactor of 5 and k = 250.

### T cell stimulation assays

Cryopreserved PBMCs were thawed, rested overnight, and plated at concentrations of 300,000 to 1 million cells per 200uL per well into a U-bottomed 96-well plate. Cells were incubated with SEB (1 ug/ml, Toxin Technology) or DMSO in the presence of monensin (2 uM, Sigma), brefeldin A (5 ug/ml, Sigma), and anti-CD107a (Biolegend). After 6 hours, cells with washed and stained with viability dyes and antibodies to CD27 and CD28 for 30 minutes at room temperature. Intracellular staining with antibodies to TNF-α, IFN-γ, Granzyme B, perforin, CD3, CD8, and CD4 (Table S3) was performed following BD Cytofix/Cytoperm Fixation/Permeabilization Kit according to manufacturer protocol (BD). For tofacitinib treatments, PBMCs were incubated with10 uM of tofacitinib (MedChemExpress) for 40 hours, after which the supernatant was removed and replaced with fresh media containing SEB, anti-CD107a, brefeldin, and monensin for a 6-hour stimulation as above. After antibody staining, cells were fixed with 2% paraformaldehyde, acquired on LSRII (BD), and analyzed by FlowJo (BD). Tofacitinib inhibition was calculated as 1- (treated/untreated), with 0 as the lower bound.

### Statistics

Normality was assessed using D’Agostino-Pearson test. Nonparametric test was used if any of the variables were non-normally distributed. Otherwise, parametric test was used. For Pearson or Spearman correlations, least squares linear regression was used to calculate the best-fitting line. Statistical comparisons were performed using Welch’s t-test, paired t-test, Mann-Whitney test, Wilcoxon matched-pairs signed rank test, and repeated measures (RM)-two-way ANOVA. Associations with RA disease characteristics were assessed using univariate linear regression. A p-values of <0.05 was used as the significance level and adjusted if multiple comparisons were performed. Statistical analyses were performed using GraphPad Prism. Lines and bars represent the mean and variability is represented by the standard error of the mean (SEM). ^*^ P < 0.05, ^**^ P < 0.01, ^***^ P < 0.001, ^****^ P < 0.0001.

### Study approval

All participants have given written informed consent. The study was approved by the Institutional Review Board at the University of Pennsylvania, Philadelphia (approval 819711, 820884) and Corporal Michael J Crescenz VA Medical Center (approval 01480).

## Data availability

All data needed to evaluate the conclusions in the paper are present in the paper or the supplementary materials. Analyses are performed using standard analysis packages. All data points are reported in the Supporting Data Values file. Further information is available from the corresponding author on request.

## Supplementary Material

**Figure S1:**
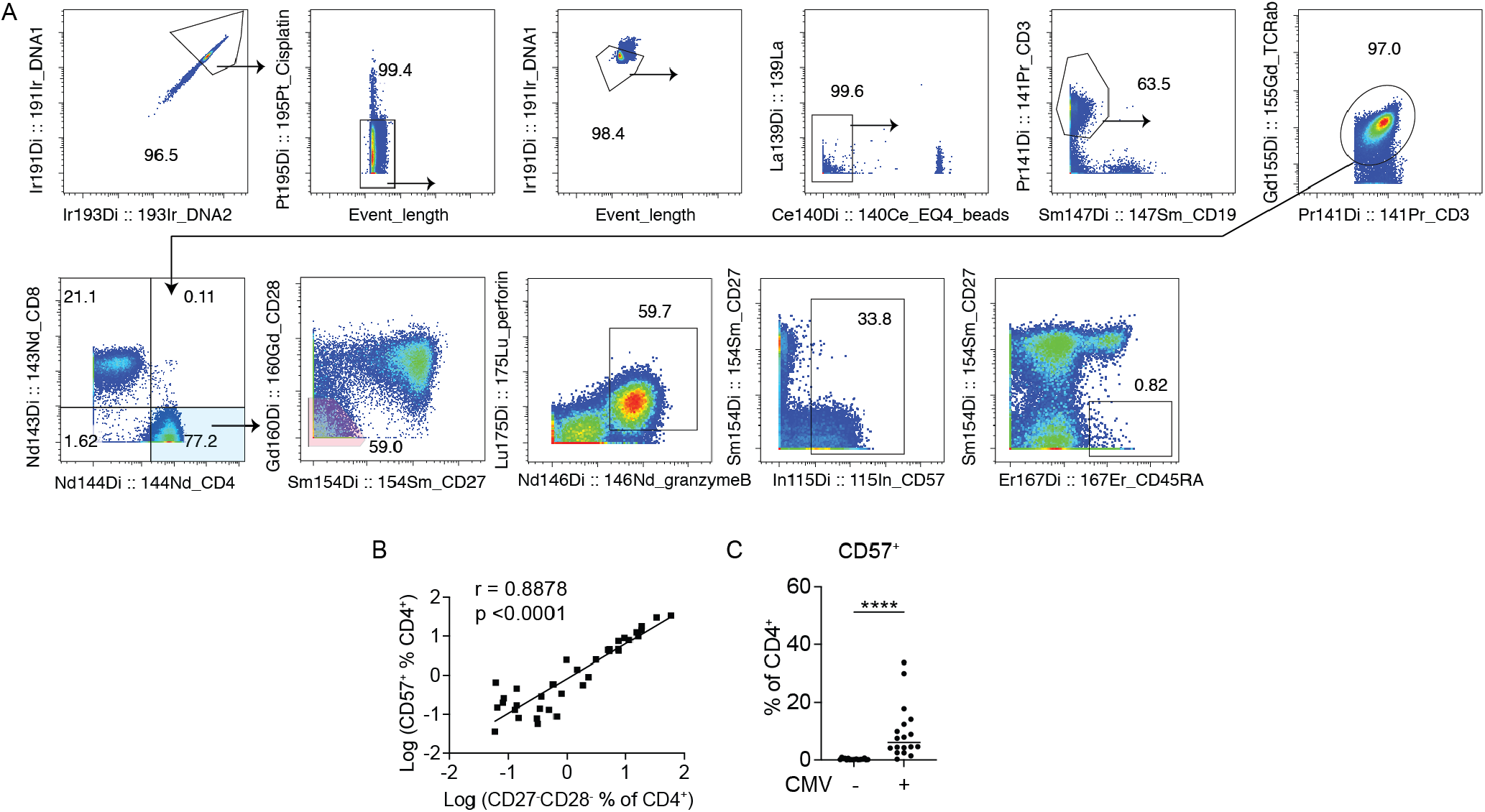
Identification of various differentiated CD4^+^ T cell subsets by mass cytometry. (A) Representative plots show the gating strategy for identifying total CD4^+^ T cells and CD27^-^CD28^-^, GzmB^+^Perf^+^, CD57^+^, CD45RA^+^CD27^-^ subsets in the PBMCs from a CMV seropositive individual. Cells exported for Spectral analyses are highlighted in blue (total CD4^+^) and red (CD27^-^CD28^-^CD4^+^). (B) The correlation between CD27^-^ CD28^-^CD4^+^ T cells and CD57^+^CD4^+^ T cells. (C) The frequency of manually gated CD57^+^ subset as a percentage of CD4^+^ T cells in CMV seronegative and seropositive donors. Each symbol represents cells from one individual. Spearman correlation was performed in (B). Mann-Whitney test was used for (C).

**Figure S2:**
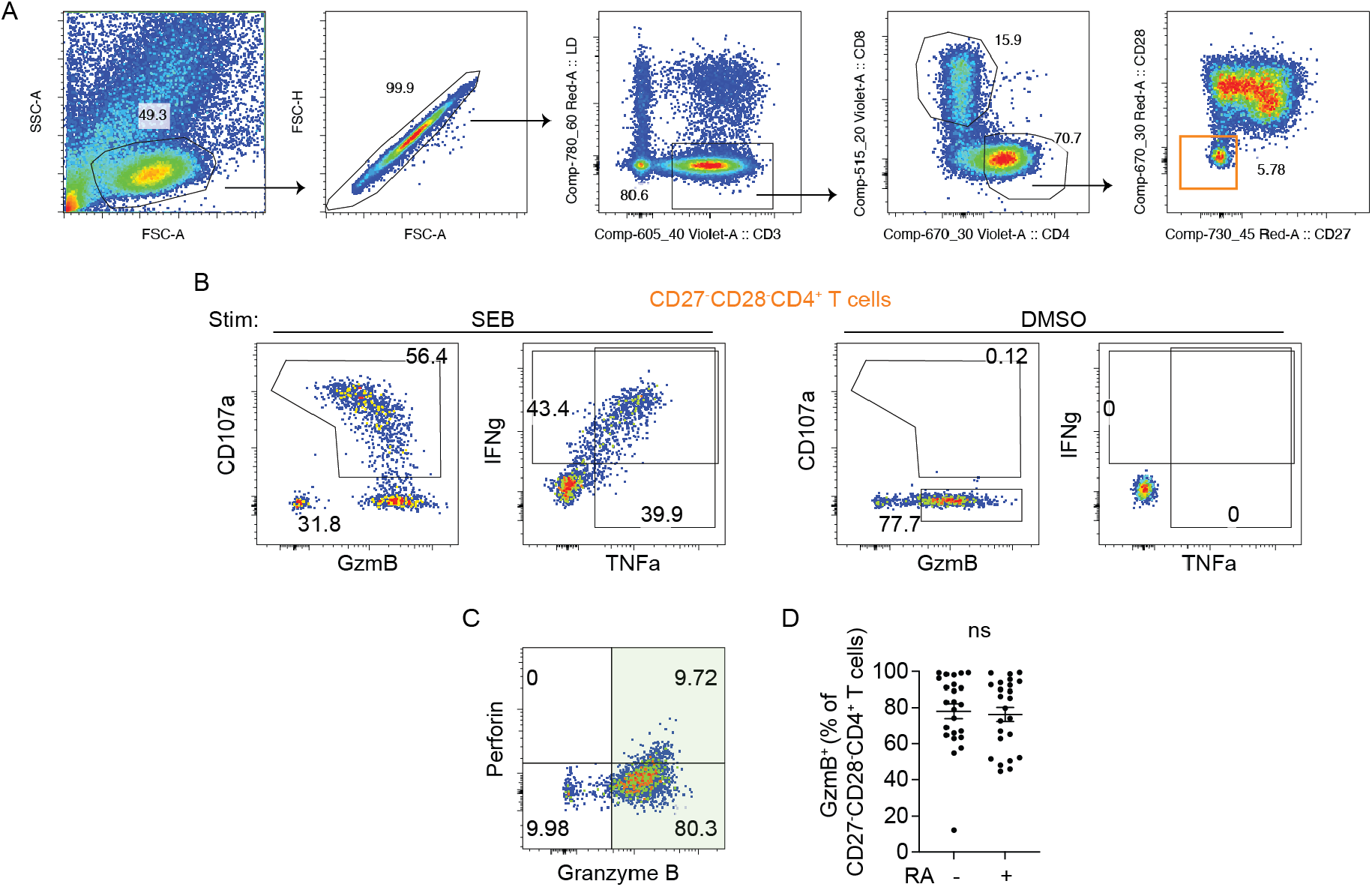
Functional responses of senescent CD4^+^ T cells to SEB stimulation. (A) FACS plots show the gating strategy for identifying CD27^-^CD28^-^CD4^+^ T cells. (B) T cell response after a 6-hour treatment with SEB or DMSO control. Plots show representative staining for TNF-α, IFN-γ, or CD107 expression within the CD27^-^ CD28^-^CD4^+^ T cell population from a CMV seropositive RA patient. (C-D) Representative plot showing perforin and GzmB expression in the DMSO-treated condition (C). The frequency of GzmB-expressing cells, highlighted in green, is summarized by disease groups (D). Mann-Whitney test was performed.

**Figure S3:**
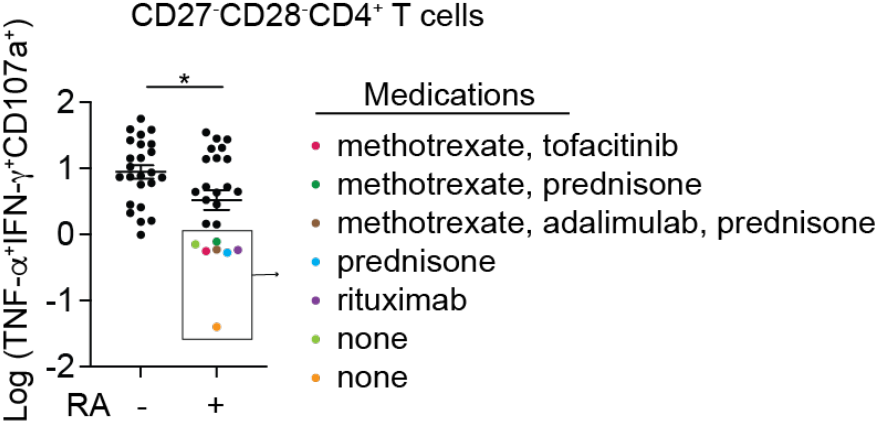
Medications taken by RA patients with severely reduced CD27^-^CD28^-^CD4^+^ T cell function. The rectangle highlights RA patients with fewer than 1% of TNF-α^+^IFN-γ ^+^CD107^+^ cells within the CD27^-^CD28^-^ CD4^+^ T cell subset. Medications taken by each RA patient are color-coded as indicated.

**Table S1:** Associations between CD27^-^CD28^-^CD4^+^ T cell functional parameters and medication use by linear regression.

|  | TNF Biologic Use | Non-TNF Biologic Use | Tofacitinib Use |
| --- | --- | --- | --- |
| Log TNF-a | -0.067 (-1.03, 0.90) | -0.43 (-1.32, 0.46) | -0.58 (-1.80, 0.65) |
| Log IFN-g | 0.067 (-1.03, 0.90) | -0.40 (-1.34, 0.53) | -0.56 (-1.82, 0.71) |
| Log CD107<sup>+</sup> | 0.049 (-0.88, 0.98) | -0.37 (-1.23, 0.50) | -0.88 (-2.06, 0.29) |
| Log TNF-a<sup>+</sup>IFN-g<sup>+</sup>CD107<sup>+</sup> | 0.20 (-0.95, 1.35) | -0.29 (-1.40, 0.80) | -0.97 (-2.45, 0.51) |

**Table S2:** List of CyTOF antibodies.

| Antigen | Metal | Clone | Vendor | Staining |
| --- | --- | --- | --- | --- |
| CD57 | 115 | HCD57 | biolegend | surface - post-cell-ID |
| CD3 | 141 | UCHT1 | BD | surface - post-cell-ID |
| CD56 | 142 | HCD56 | Biolegend | surface - post-cell-ID |
| CD8 | 143 | SK1 | Biolegend | surface - post-cell-ID |
| CD4 | 144 | SK3 | Biolegend | surface - post-cell-ID |
| CD16 | 145 | B73.1 | Biolegend | surface - post-cell-ID |
| granzyme B | 146 | CLB-GB11 | eBioscience | intracellular |
| CD19 | 147 | HIB19 | Biolegend | surface - post-cell-ID |
| HLA-DR | 148 | L243 | Biolegend | surface - post-cell-ID |
| CD127 | 150 | A019D5 | 351302 | surface - pre cell-ID |
| CD38 | 151 | HIT2 | Biolegend | surface - post-cell-ID |
| CTLA4 | 152 | BNI3 | BD | intracellular |
| CCR2 | 153 | K036C2 | Biolegend | surface - pre cell-ID |
| CD27 | 154 | LG.7F9 | eBioscience | surface - pre cell-ID |
| TCRab | 155 | T10B9.1A-31 | BD | surface - pre cell-ID |
| CXCR3 | 156 | G025H7 | Biolegend | surface - pre cell-ID |
| PD-1 | 158 | EH12.2H7 | Biolegend | surface - post-cell-ID |
| CD28 | 160 | CD28.2 | Biolegend | surface - post-cell-ID |
| CD11b | 162 | ICRF44 | Biolegend | surface - post-cell-ID |
| ICOS | 163 | C398.4A | Biolegend | surface - post-cell-ID |
| Ki67 | 164 | B56 | BD | intracellular |
| Foxp3 | 165 | PCH101 | eBiosciences | intracellular |
| TCF1 | 166 | C63D9 | Cell Signaling Technology | intracellular |
| CD45RA | 167 | HI100 | eBio | surface - post-cell-ID |
| SIRPa | 168 | 15-414 | Biolegend | surface - post-cell-ID |
| granulysin | 169 | DH2 | Biolegend | intracellular |
| CXCR5 | 170 | RF8B2 | BD | surface - post-cell-ID |
| CD94 | 171 | DX22 | Biolegend | surface - post-cell-ID |
| 2B4 | 172 | C1.7 | Biolegend | surface - post-cell-ID |
| CCR6 | 173 | G034E3 | G034E3 | surface - pre cell-ID |
| CD39 | 174 | ebioA1 | eBiosciences | surface - post-cell-ID |
| perforin | 175 | dG9 | Biolegend | intracellular |
| CD95 | 176 | DX2 | Biolegend | surface - post-cell-ID |

**Table S3:** List of flow cytometry antibodies.

| Antigen | Fluorochrome | Clone | Vendor | Catalog # |
| --- | --- | --- | --- | --- |
| CD107a | PE-TxR | H4A3 | Biolegend | 328646 |
| Live/Dead | Near-IR | NA | Invitrogen | L10119 |
| CD27 | AF700 | O323 | Biolegend | 302814 |
| CD28 | APC | CD28.2 | Biolegend | 302911 |
| CD45RO | BV650 | UCHL1 | Biolegend | 304231 |
| CD3 | BV605 | OKT3 | Biolegend | 317322 |
| CD4 | BV510 | SK3 | Biolegend | 344634 |
| CD8 | PE-Cy5 | HIT8a | Biolegend | 300910 |
| Granzyme B | PE-Cy7 | QA18A28 | Biolegend | 396410 |
| Perforin | PE | B-D48 | Biolegend | 353304 |
| TNF-α | BV785 | MAb11 | Biolegend | 502948 |
| IFN-γ | FITC | 4S.B3 | Biolegend | 502506 |

NA: not applicable

## References

1. Maeda T, Yamada H, Nagamine R, Shuto T, Nakashima Y, Hirata G, et al. Involvement of CD4+,CD57+ T cells in the disease activity of rheumatoid arthritis. Arthritis Rheum. 2002;46(2):379–379.

2. Goronzy JJ, Matteson EL, Fulbright JW, Warrington KJ, Chang-Miller A, Hunder GG, et al. Prognostic markers of radiographic progression in early rheumatoid arthritis. Arthritis Rheum. 2004;50(1):43–43.

3. Pawlik A, Ostanek L, Brzosko I, Brzosko M, Masiuk M, Machalinski B, et al. The expansion of CD4+CD28-T cells in patients with rheumatoid arthritis. Arthritis Res Ther. 2003;5(4):R210–3.

4. Winchester R, Giles JT, Nativ S, Downer K, Zhang HZ, Bag-Ozbek A, et al. Association of Elevations of Specific T Cell and Monocyte Subpopulations in Rheumatoid Arthritis With Subclinical Coronary Artery Atherosclerosis. Arthritis & rheumatology. 2016;68(1):92–92.

5. Gerli R, Schillaci G, Giordano A, Bocci EB, Bistoni O, Vaudo G, et al. CD4+CD28-T lymphocytes contribute to early atherosclerotic damage in rheumatoid arthritis patients. Circulation. 2004;109(22):2744–2744.

6. Abo T, and Balch CM. A differentiation antigen of human NK and K cells identified by a monoclonal antibody (HNK-1). J Immunol. 1981;127(3):1024–1024.

7. Laphanuwat P, Gomes DCO, and Akbar AN. Senescent T cells: Beneficial and detrimental roles. Immunol Rev. 2023;316(1):160–160.

8. Cenerenti M, Saillard M, Romero P, and Jandus C. The Era of Cytotoxic CD4 T Cells. Frontiers in immunology. 2022;13:867189.

9. van Leeuwen EM, Remmerswaal EB, Vossen MT, Rowshani AT, Wertheim-van Dillen PM, van Lier RA, et al. Emergence of a CD4+CD28-granzyme B+, cytomegalovirus-specific T cell subset after recovery of primary cytomegalovirus infection. J Immunol. 2004;173(3):1834–1834.

10. Fletcher JM, Vukmanovic-Stejic M, Dunne PJ, Birch KE, Cook JE, Jackson SE, et al. Cytomegalovirus-specific CD4+ T cells in healthy carriers are continuously driven to replicative exhaustion. J Immunol. 2005;175(12):8218–8218.

11. Brenchley JM, Karandikar NJ, Betts MR, Ambrozak DR, Hill BJ, Crotty LE, et al. Expression of CD57 defines replicative senescence and antigen-induced apoptotic death of CD8+ T cells. Blood. 2003;101(7):2711–2711.

12. Palmer BE, Blyveis N, Fontenot AP, and Wilson CC. Functional and phenotypic characterization of CD57+CD4+ T cells and their association with HIV-1-induced T cell dysfunction. J Immunol. 2005;175(12):8415–8415.

13. Brown DM, Lee S, Garcia-Hernandez Mde L, and Swain SL. Multifunctional CD4 cells expressing gamma interferon and perforin mediate protection against lethal influenza virus infection. J Virol. 2012;86(12):6792–6792.

14. Wilkinson TM, Li CK, Chui CS, Huang AK, Perkins M, Liebner JC, et al. Preexisting influenza-specific CD4+ T cells correlate with disease protection against influenza challenge in humans. Nat Med. 2012;18(2):274–274.

15. Oh DY, Kwek SS, Raju SS, Li T, McCarthy E, Chow E, et al. Intratumoral CD4(+) T Cells Mediate Anti-tumor Cytotoxicity in Human Bladder Cancer. Cell. 2020;181(7):1612–1612e13.

16. Quezada SA, Simpson TR, Peggs KS, Merghoub T, Vider J, Fan X, et al. Tumor-reactive CD4(+) T cells develop cytotoxic activity and eradicate large established melanoma after transfer into lymphopenic hosts. J Exp Med. 2010;207(3):637–637.

17. Zhang L, Yu X, Zheng L, Zhang Y, Li Y, Fang Q, et al. Lineage tracking reveals dynamic relationships of T cells in colorectal cancer. Nature. 2018;564(7735):268–268.

18. Cachot A, Bilous M, Liu YC, Li X, Saillard M, Cenerenti M, et al. Tumor-specific cytolytic CD4 T cells mediate immunity against human cancer. Sci Adv. 2021;7(9).

19. Hashimoto K, Kouno T, Ikawa T, Hayatsu N, Miyajima Y, Yabukami H, et al. Single-cell transcriptomics reveals expansion of cytotoxic CD4 T cells in supercentenarians. Proc Natl Acad Sci U S A. 2019;116(48):24242–24242.

20. Finck R, Simonds EF, Jager A, Krishnaswamy S, Sachs K, Fantl W, et al. Normalization of mass cytometry data with bead standards. Cytometry A. 2013;83(5):483–483.

21. Ashhurst TM, Marsh-Wakefield F, Putri GH, Spiteri AG, Shinko D, Read MN, et al. Integration, exploration, and analysis of high-dimensional single-cell cytometry data using Spectre. Cytometry A. 2021.

22. Winter JR, Jackson C, Lewis JE, Taylor GS, Thomas OG, and Stagg HR. Predictors of Epstein-Barr virus serostatus and implications for vaccine policy: A systematic review of the literature. J Glob Health. 2020;10(1):010404.

23. Bradley H, Markowitz LE, Gibson T, and McQuillan GM. Seroprevalence of herpes simplex virus types 1 and 2--United States, 1999-2010. J Infect Dis. 2014;209(3):325–325.

24. Alamino VA, Onofrio LI, Acosta CDV, Ferrero PV, Zacca ER, Cadile, II, et al. Tofacitinib treatment of rheumatoid arthritis increases senescent T cell frequency in patients and limits T cell function in vitro. Eur J Immunol. 2023;53(8):e2250353.

25. Yan Q, Chen W, Song H, Long X, Zhang Z, Tang X, et al. Tofacitinib Ameliorates Lupus Through Suppression of T Cell Activation Mediated by TGF-Beta Type I Receptor. Frontiers in immunology. 2021;12:675542.

26. Staras SA, Dollard SC, Radford KW, Flanders WD, Pass RF, and Cannon MJ. Seroprevalence of cytomegalovirus infection in the United States, 1988-1994. Clin Infect Dis. 2006;43(9):1143–1143.

27. Fowler K, Mucha J, Neumann M, Lewandowski W, Kaczanowska M, Grys M, et al. A systematic literature review of the global seroprevalence of cytomegalovirus: possible implications for treatment, screening, and vaccine development. BMC Public Health. 2022;22(1):1659.

28. Dollard SC, Staras SA, Amin MM, Schmid DS, and Cannon MJ. National prevalence estimates for cytomegalovirus IgM and IgG avidity and association between high IgM antibody titer and low IgG avidity. Clin Vaccine Immunol. 2011;18(11):1895–1895.

29. Pierer M, Rothe K, Quandt D, Schulz A, Rossol M, Scholz R, et al. Association of anticytomegalovirus seropositivity with more severe joint destruction and more frequent joint surgery in rheumatoid arthritis. Arthritis Rheum. 2012;64(6):1740–1740.

30. Hooper M, Kallas EG, Coffin D, Campbell D, Evans TG, and Looney RJ. Cytomegalovirus seropositivity is associated with the expansion of CD4+CD28- and CD8+CD28-T cells in rheumatoid arthritis. J Rheumatol. 1999;26(7):1452–1452.

31. Ytterberg SR, Bhatt DL, Mikuls TR, Koch GG, Fleischmann R, Rivas JL, et al. Cardiovascular and Cancer Risk with Tofacitinib in Rheumatoid Arthritis. N Engl J Med. 2022;386(4):316–316.

32. Curtis JR, Yamaoka K, Chen YH, Bhatt DL, Gunay LM, Sugiyama N, et al. Malignancy risk with tofacitinib versus TNF inhibitors in rheumatoid arthritis: results from the open-label, randomised controlled ORAL Surveillance trial. Ann Rheum Dis. 2023;82(3):331–331.

33. Mehta B, Pedro S, Ozen G, Kalil A, Wolfe F, Mikuls T, et al. Serious infection risk in rheumatoid arthritis compared with non-inflammatory rheumatic and musculoskeletal diseases: a US national cohort study. RMD Open. 2019;5(1):e000935.

34. Winthrop KL, Yamanaka H, Valdez H, Mortensen E, Chew R, Krishnaswami S, et al. Herpes zoster and tofacitinib therapy in patients with rheumatoid arthritis. Arthritis & rheumatology. 2014;66(10):2675–2675.

35. Balanescu AR, Citera G, Pascual-Ramos V, Bhatt DL, Connell CA, Gold D, et al. Infections in patients with rheumatoid arthritis receiving tofacitinib versus tumour necrosis factor inhibitors: results from the open-label, randomised controlled ORAL Surveillance trial. Ann Rheum Dis. 2022;81(11):1491–1491.

36. Ashhurst TM, Marsh-Wakefield F, Putri GH, Spiteri AG, Shinko D, Read MN, et al. Integration, exploration, and analysis of high-dimensional single-cell cytometry data using Spectre. Cytometry A. 2022;101(3):237–237.

37. Levine JH, Simonds EF, Bendall SC, Davis KL, Amir el AD, Tadmor MD, et al. Data-Driven Phenotypic Dissection of AML Reveals Progenitor-like Cells that Correlate with Prognosis. Cell. 2015;162(1):184–97.

38. Becht E, McInnes L, Healy J, Dutertre CA, Kwok IWH, Ng LG, et al. Dimensionality reduction for visualizing single-cell data using UMAP. Nat Biotechnol. 2018.

